# *De Novo* Genes are “Frozen Accidents” which Escaped Rapid Turnover of Pervasively Transcribed ORFs

**DOI:** 10.1101/166827

**Authors:** Jonathan Schmitz, Kristian Ullrich, Erich Bornberg-Bauer

## Abstract

A recent surge of studies suggested that many novel genes arise *de novo* from previously non-coding DNA and not by duplication. However, since most studies concentrated on longer evolutionary time scales and rarely considered protein structural properties, it remains unclear how these properties are shaped by evolution, depend on genetic mechanisms and influence gene survival. Here we compare open reading frames (ORFs) from high coverage transcriptomes from mouse and another four mammals covering 160 million years of evolution. We find that novel ORFs pervasively emerge from intergenic and intronic regions but are rapidly lost again while relatively fewer arise from duplications but are retained over much longer times. Surprisingly, disorder and other protein properties of young ORFs do not change with gene age. Only length and nucleotide composition change, probably to avoid aggregation. Thus de novo genes resemble frozen accidents of randomly emerged ORFs which survived initial purging, likely because they are functional.

## Background

Sequencing of hundreds of genomes showed that all genomes contain some 10-30% protein-coding genes (CDS) without computationally detectable similarity to proteins from other genomes (Tautz et al. 2011; Khalturin et al. 2009). Previously, such “orphan” genes were thought to be novel genes which arise exclusively through the fast divergence of duplicated genes (Ohno 1970; Zhang 2003; Long et al. 2003; Domazet-Loso et al. 2003). However, accumulating evidence suggests frequent *de novo* emergence (Wissler et al. 2013; Wu et al. 2011; Donoghue et al. 2011), i.e. the evolution of a formerly non-coding DNA sequence towards a CDS (Begun et al. 2007; Carvunis et al. 2012). It is, however, unclear if ORFs which arose from intergenic DNA and have not experienced selection pressure to form a structured and functional protein, are in general deleterious and pose a fitness burden to the organism, if they are mostly neutral or how likely they can be beneficial and become fixed.

On the one hand, de novo proteins could easily misfold or aggregate (Monsellier et al. 2007; Geiler-Samerotte et al. 2011) and therefore be toxic to the cell. Proteins seem to have evolved to exhibit a certain amount of stability with much more stability leading to aggregation and too little stability to misfolding (DePristo et al. 2005). In this framework, it seems unlikely that proteins encoded by intergenic ORFs would exhibit exactly the correct amount of stability to avoid aggregation or misfolding. On the other hand, it is also concievable that de novo proteins might fold, at least to some extent, be not detrimential and soon become exposed to selection. Supporting lines of evidence include, among others, early theoretical studies (Ptitsyn 1985), a recent computational study suggesting that proteins should not aggregate for varying GC-contents (Ángyán et al. 2012), and the observation that chaperones often assist in the correct folding of proteins (Saibil 2013).

But, even in the absence of harmful effects, newly transcribed intergenic ORFs would need to code for functional proteins to avoid being purged by drift. Indeed, intrinsically disordered, i.e., unstructured proteins were, until recently, to be harmful, but are now known to have many important functions (Tompa 2011) and sometimes disorder and the transition to order is even essential (Wright et al. 2015; Bellay et al. 2011). Accordingly, disorder in fixed, i.e., non-detrimental *de novo* genes may be either a selected for trait or a concomittant trait which helped, under certain circumstances, acquire a function – for example binding – for which the protein can become beneficial. While an earlier (Carvunis et al. 2012) study suggested disorder increased with age of *de novo* genes, others reported the opposite trend (Zhao et al. 2014; Bornberg-Bauer, Jonathan Schmitz, et al. 2015; Wilson et al. 2017). Conceivable, results may be affected by parameter settings in disorder prediction programmes and the fact that genes not always emerge *de novo* in their entirety but that domains may emerge *de novo* by ORF extentions or exonisation of intronic regions (Bornberg-Bauer, Jonathan Schmitz, et al. 2015; Bornberg-Bauer and Albà 2013).

Apart from binding, proteins from random, artificial libraries and without prior selection also showed to be beneficial in vitro (Keefe et al. 2001) and in vivo in *Escherichia coli* (Neme, Amador, et al. 2017). Presence and nature of their functional roles remain unclear, but aside binding, weak enzymatic activities may arise too (Hollfelder et al. 2000).

Considering these inconsistencies and ambiguities about the functional potential of *de novo* genes evokes questions regarding selection pressure and retention rate during their early evolution. Previous analyses found that *de novo* (Zhao et al. 2014; J.-Y. Chen et al. 2015) and also all novel genes (Palmieri et al. 2014) accumulate fewer mutations than intergenic regions. But even though selection appears to act on novel genes, younger novel genes have a higher loss probability compared to older novel genes and to more established genes in general (Palmieri et al. 2014; Wissler et al. 2013). Novel genes (S. Chen et al. 2010) and *de novo* genes in particular (Reinhardt et al. 2013; Gubala et al. 2017) often contribute significantly to an organism’s fitness but when, why and how strongly selection sets in and novel genes become fixed remains unclear too.

Most early studies on the origin of novel genes used *Drosophila* genomes comparing genes predicted by standard methods and focused on confirming the *de novo* emergence of novel genes (Long et al. 2003; Levine et al. 2006; Begun et al. 2007). Confirmed by studies on Mammalia (Knowles et al. 2009; Wu et al. 2011; J.-Y. Chen et al. 2015; Guerzoni et al. 2016), they showed that *de novo* genes can emerge when intergenic ORFs become transcribed. So far, only few and mostly ancillary results considered possible cross-dependencies between genetic mechanisms and protein properties but without any consensus (Carvunis et al. 2012; Ángyán et al. 2012; Abrusán 2013; Wilson et al. 2017). Also, the raw material for de novo emergence, i.e. ORFs from non-genic transcripts, is largely understudied. This is surprising considering that other lines of research revealed a pervasively transcribed genome (Neme and Tautz 2016; Kapranov et al. 2012) and the exonisation of intergenic DNA and of intronic regions in particular (Singer et al. 2004; Krull et al. 2005; Jürgen Schmitz et al. 2011). Novel transcripts abound, e.g., in mouse (Neme and Tautz 2016), but properties of ORFs contained and possible functions of proteins encoded remain unclear.

Here we compare a high-coverage mouse (*Mus musculus*) transcriptome with four other mammalian transcriptome data sets spanning an evolutionary range of 160 mya (million years ago) to assess conservation and age of novel ORFs. Subsequently, we computationally analyse sequence properties of ORFs and structural properties of the encoded proteins to evaluate the coding potential of transcribed intergenic ORFs and the evolutionary changes of these properties after gene emergence. Finally, we compare properties and emergence mechanisms of *de novo* ORFs with ORFs from randomly generated sequences and ORFs created by other mechanisms.

## Results & Discussion

### ORF annotation status varies with ORF age

Here we used 5 high-coverage transcriptomes from mouse and 4 other species that we assembled ensuring comparable data quality (see Methods section for details). We first analysed the genomic position of all ORFs present on assembled mouse transcripts, to quantify and analyse the raw materials of *de novo* gene emergence (see Methods). For our first analyses, we only considered the longest plus-strand ORFs (l+ORFs) found on each transcript (Figure 1, Table 2). l+ORFs are usually seen as the only biologically relevant transcript (Kozak 1999; Mouilleron et al. 2016). Interestingly, most of the l+ORFs analysed here are either mouse-specific or conserved across all analysed species, i.e., including opossum. These latter l+ORFs emerged at least 160 mya, mostly (79%) correspond to genes which are already annotated in the mouse genome and thus confirm that many established genes in eukaryotic genomes are very old.

**Figure 1.**
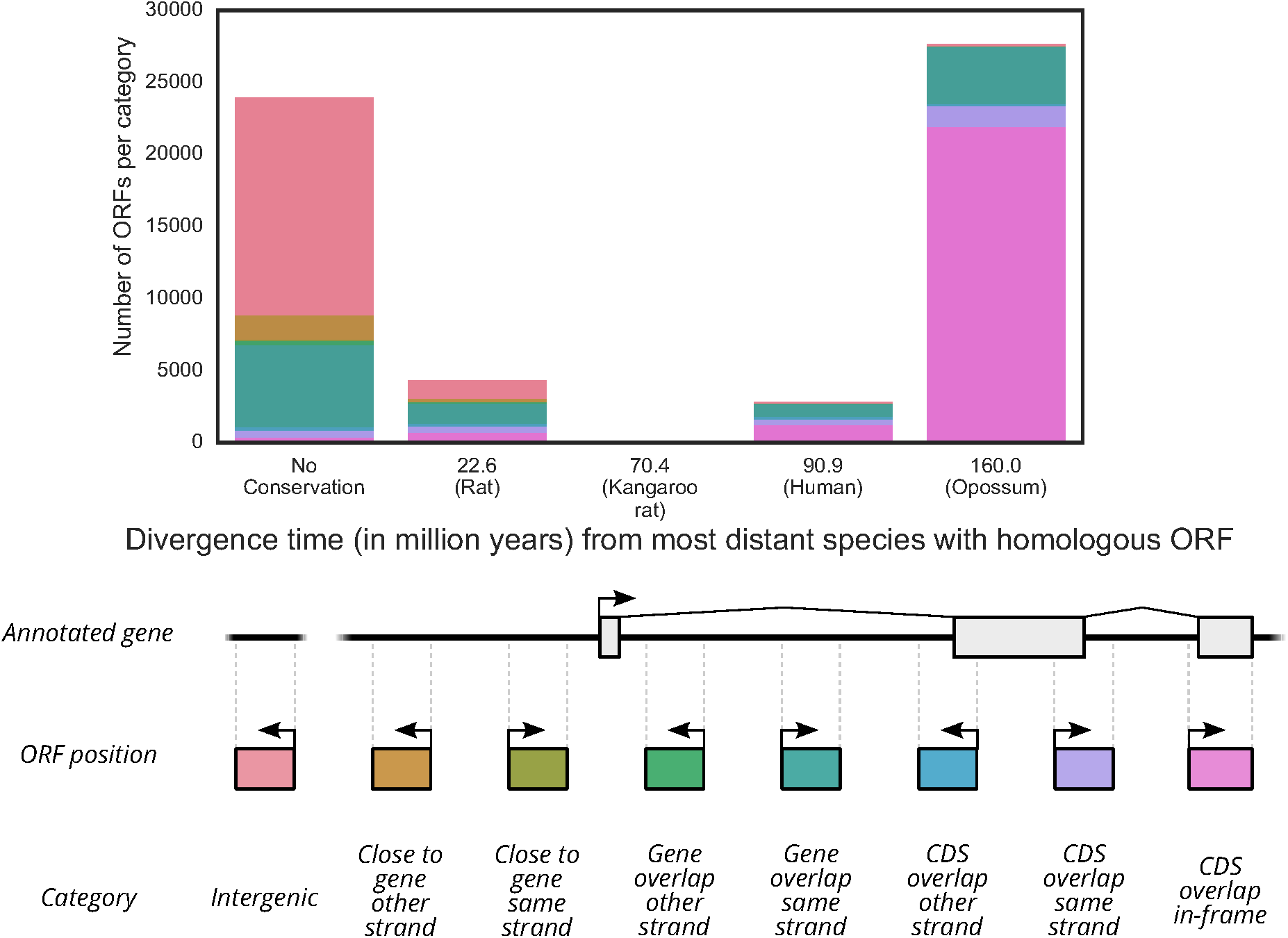
**a: Number of ORFs per age class and annotation category.** Only the longest plus-strand ORF from each transcript is depicted here. The “Mouse specific” class contains ORFs for which no homologs could be found in any of the other species’ transcriptomes. Each ORF is only counted for the most distant species with a similar ORF, i.e. if an ORF has homologs in rat, kangaroo rat, and human, it would only be counted in the “Human” age class, **b: Pictogram describing the different annotation categories.** Genes within 5kb of ORFs were considered to be close. The classes are hierarchical, i.e. an ORF overlapping with CDS also overlaps with the gene, but only the rightmost class is considered for each ORFs. Introns are seen as parts of genes here, so “Intergenic” ORFs can not overlap witn mtrons or genes.

Most mouse-specific 1+ORFs, on the other hand, reside either in intergenic regions (15,163, 63%), relative to the annotated mouse genes, or inside introns of annotated genes, on the same strand (5,656, 24%). Interestingly, only few mouse-specific 1+ORFs are located e.g. on the opposite strand of established genes. Accordingly, many fully processed mRNAs stem from introns of established genes. This finding matches with the fact that introns of established genes are transcriptionally active regions and earlier incidential observations that novel genes can emerge in introns (Gubala et al. 2017). The high number of intergenic and intronic mouse-specific 1+ORFs also fits expectation that many mouse-specific ORFs are found on spurious and low-frequency transcripts (Neme and Tautz 2016) and as such represent currently emerged ORFs and transcripts. However, there seems to be a high loss-rate of these ORFs too, as only relatively few (28%) of them are found in the other species. Probably, as a consequence of this high turnover rate, many of these ORFs will lose transcription again. An additional factor contributing to the high number of unannotated mouse-specific 1+ORFs could be that mouse-specific genes are more difficult to annotate than older genes, as they can not be annotated based on sequence homology due to their nature as novel genes (J. F. Schmitz et al. 2017).

We next analysed how all other ORFs found on transcripts fare over evolutionary long time frames because more than one ORF per transcript might be functional (Mouilleron et al. 2016) and small ORFs (smORFs) can also be conserved over long evolutionary distances (Ladoukakis et al. 2011; Couso 2015; Mackowiak et al. 2015) and functional (Galindo et al. 2007). We find a higher fraction of alternative ORFs originating from introns of annotated genes than for l+ORFs, particularly on the opposite strand relative to the annotated gene. Equal numbers of ORFs were found on the forward and on the reverse strand of transcripts (Supplementary Figures S 1 and S 2; Table 2). Interestingly, also some (ca. 40,000) ORFs overlapping with established genes on the opposite strand are conserved since 160mya. This finding underlines a potential functional role, e.g. as lncRNAs (long non-coding RNAs) (Heinen et al. 2009; Ruiz-Orera et al. 2014; Bornberg-Bauer and Albà 2013), as such patterns of conservation are unlikely to occur by chance as non-functional ORFs would be expected to pseudogenise rapidly.

### Only length and nucleotide composition correlate strongly with ORF age

Next, we computationally analysed the nucleotide sequence properties of ORFs as well as the structural properties of the encoded proteins and classified the ORFs according to their age by determining the most recent common ancestor of all occurrences with a detectable homology. Again, we first restricted our analysis to only include the l+ORF (Figure 2 and Table 1). We analysed nucleotide composition properties such as hexamer score as well as a number of predicted structural properties including disorder fraction, aggregation propensity, hydrophobic cluster coverage, and mean hydrophobicity (see Methods section for details). Hexamer score and sequence length were the only sequence properties to correlate noticeably with ORF age.

**Figure 2.**
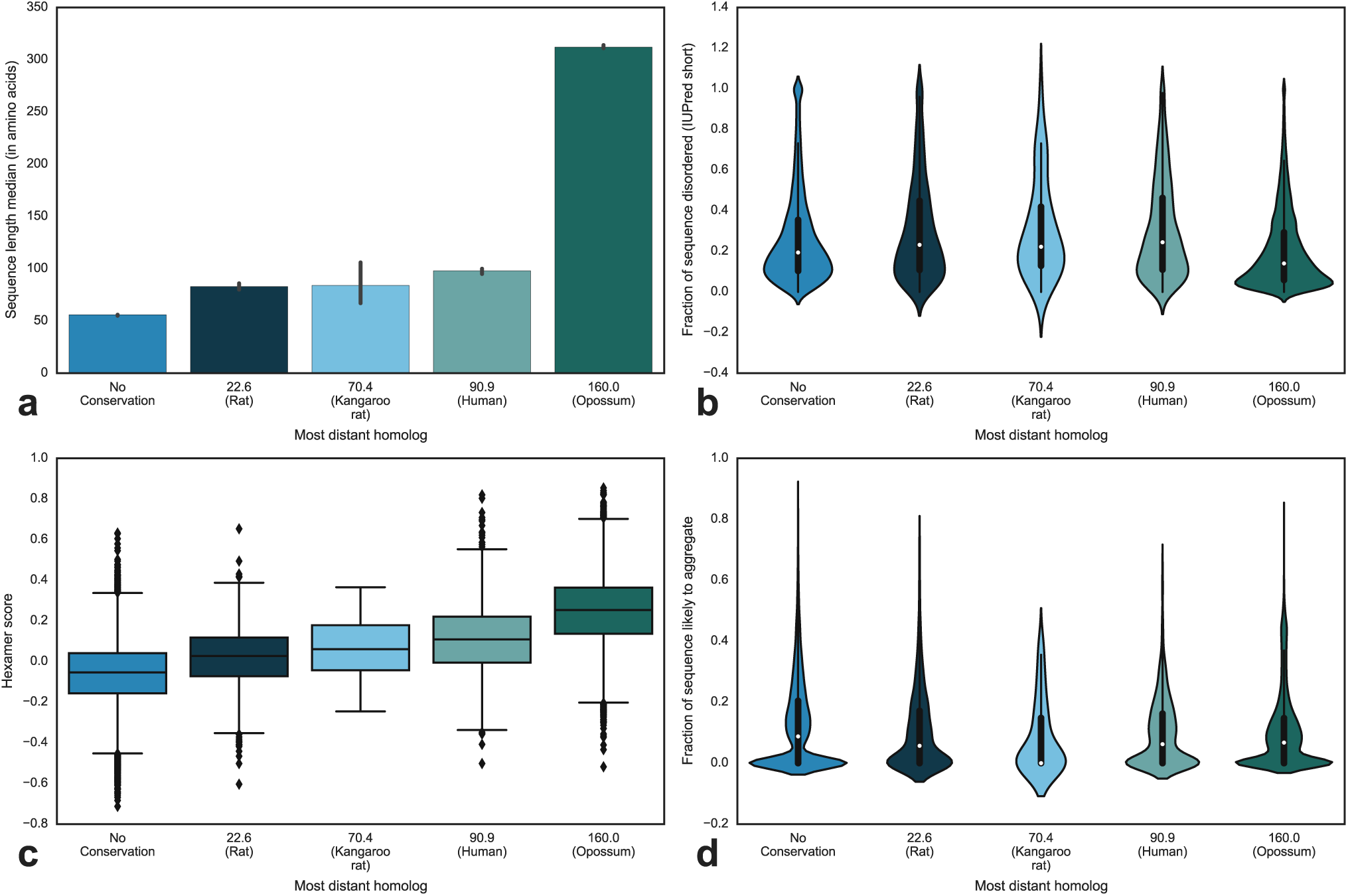
Comparison of ORF sequence properties across age classes. Only the longest plus-strand ORF per transcript was considered here. Age classes are defined by the most distant species with homologous sequences. Data are shown as violin plots. Overlaid are box plots with a white dot denoting the median value. **a**: Median sequence length. The black line indicates the 95% confidence interval based on bootstrapping. **b**: Fraction of each sequence found to be disordered using the IUPred short algorithm. **c**: Hexamer score of each sequence calculated using CPAT (Wang et al. 2013). **d**: Fraction of each sequence predicted to aggregate, calculated using TANGO (Fernandez-Escamilla et al. 2004; Linding et al. 2004).

**Table 1.**
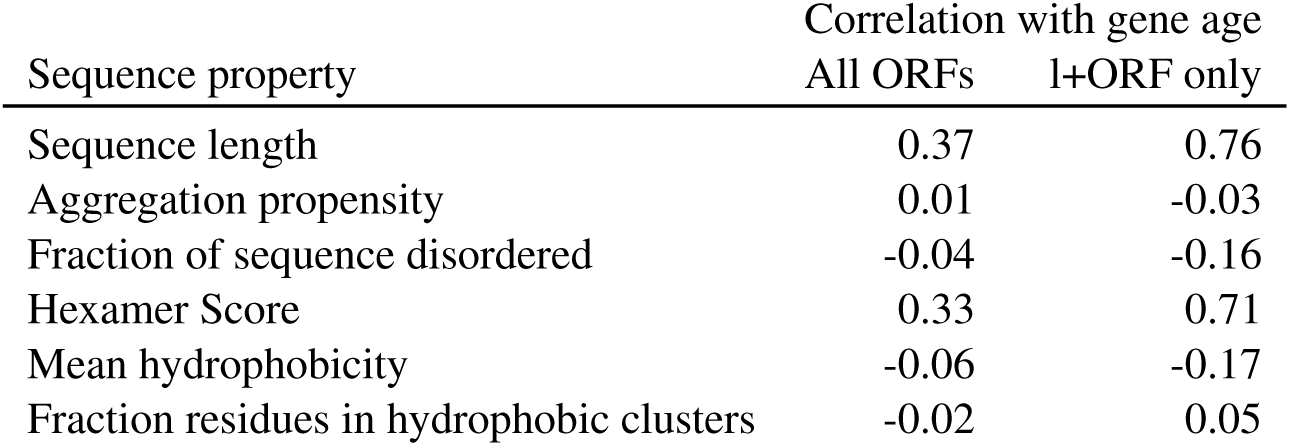
Spearman rank correlation coefficients of age with the analysed sequence properties. All correlations are highly significant with p-values << 0.001 after FDR correction

| Sequence property | Correlation with gene age |  |
| --- | --- | --- |
|  | All ORFs | l+ORF only |
| Sequence length | 0.37 | 0.76 |
| Aggregation propensity | 0.01 | -0.03 |
| Fraction of sequence disordered | -0.04 | -0.16 |
| Hexamer Score | 0.33 | 0.71 |
| Mean hydrophobicity | -0.06 | -0.17 |
| Fraction residues in hydrophobic clusters | -0.02 | 0.05 |

**Table 2.**
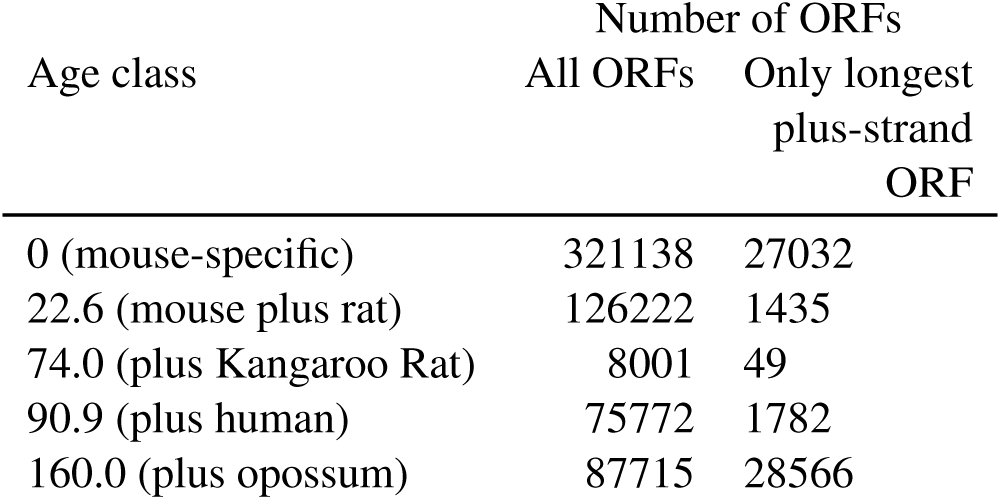
Numbers of ORFs found per age class. Number of ORFs

In a second step, we again included all ORFs from each transcript. Here, we find a pattern similar to the one for l+ORFs (Table 1 and Supplementary Figure S 3), but with lower correlation coefficients. This difference in correlation strengths likely reflects that the l+ORF is most strongly shaped by adaptation.

The low correlation of the predicted structural properties with ORF age (Table 1) shows that novel genes in general do not strongly change amino acid composition after their initial emergence and fixation in the genome, at least not in the analysed time frame. Accordingly, the overall structural composition of emerging and intergenic ORFs might already be sufficient to avoid detrimental functions and ORFs do not change their structural characteristics drastically after emergence. Still, a gradual adaptation of general amino acid composition might take place over even longer time periods and the evolutionary forces acting on longer genes might be different (Bornberg-Bauer, Jonathan Schmitz, et al. 2015). However, some nucleotide sequence properties such as length and hexamer score do change in the time frame analysed here. This finding suggests that at least ORF length expansion and hexamer usage are generally favored by selection or, alternatively, a by-product of hitherto undetected evolutionary forces acting on protein-coding sequences in general.

### Young ORFs have similar properties to randomly generated ORFs

Next, we compared the predicted sequence properties of randomly generated ORFs and the predicted structural properties of the proteins encoded by them with those of the actually transcribed, unconserved ORFs to detect potential biases in the process by which ORFs gain transcription (Figure 3; see also Supplementary Figure S 4). Such biases would suggest that not all intergenic ORFs have the potential to become protein-coding genes. For this purpose we generated random nucleotide sequences based on the hexamer composition of CDS, genic, and intergenic mouse DNA, respectively (see Materials & Methods for details). This approach was chosen to identify the influence of the overall genomic nucleotide composition on the properties of occurring ORFs. Using this method, we analysed ORFs that can occur incidentally in the genome and allow even for ORFs that would be selected against if they were to occur in an actual organism. Our approach factored out direct selection on ORF properties and simply asks what kinds of ORFs (and encoded proteins) the genomic nucleotide composition taken together with the genetic code produces, without selection playing a role.

**Figure 3.**
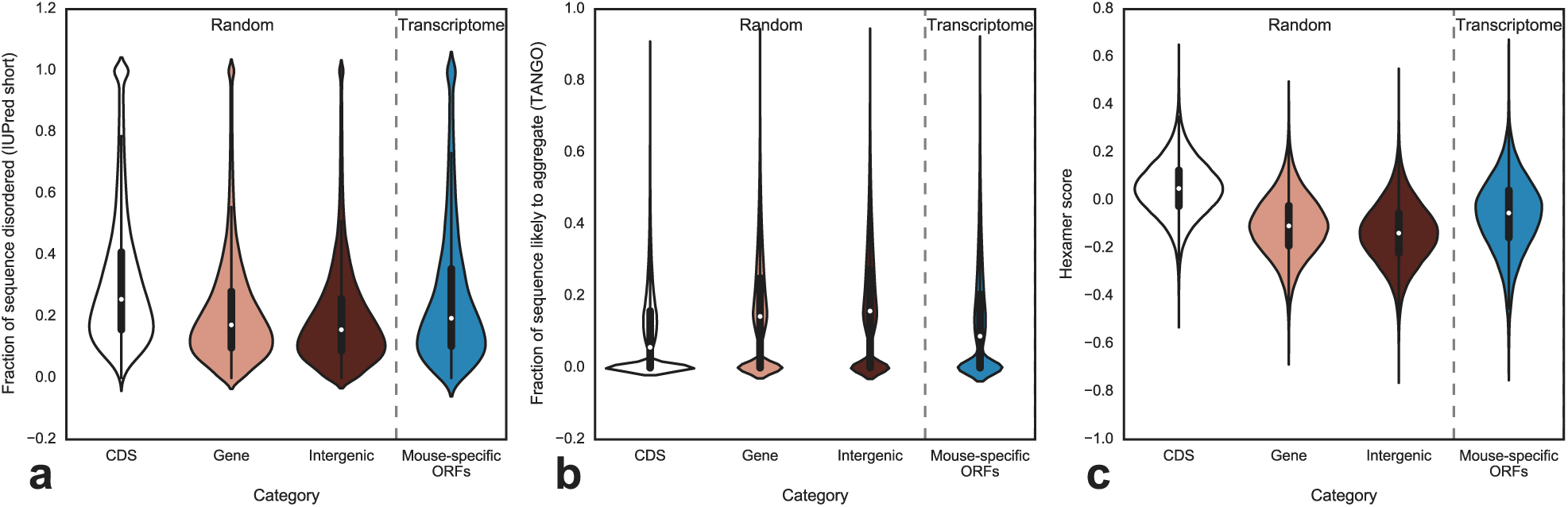
**Comparison of unconserved ORF sequences to randomly generated sequences**. Random sequences were generated based on the hexamer content of CDS, genic and intergenic mouse sequences. Overlaid over violin plots indicating the density distribution are box plots with a white dot denoting the median value. **a**: Comparison of the disordered sequence fraction. **b**: Fraction of each sequence likely to aggregate, calculated using TANGO (Fernandez-Escamilla et al. 2004; Linding et al. 2004). **c**: Sequence hexamer score.

In this analysis, differences can be observed for some properties between the ORFs found on the different random sequence classes. The ORFs represent sequences found between start and stop codons in pseudo-random nucleotide sequences described above. Specifically, random ORFs generated from the CDS hexamer distribution exhibit a higher GC-content, a higher disordered fraction, a higher hexamer score, less aggregation propensity, and a lower coverage of hydrophobic clusters compared to the ORFs generated from genic and intergenic hexamer distributions. The ORFs randomly generated from the intergenic and genic hexamer distributions are highly similar to each other for most properties. The observed trends could be caused by a combination of the higher general GC-content of coding sequences (Supplementary Figure S 4) and higher likelihood of CDS-kmers to code for disorder-promoting amino acids. This higher likelihood would have an effect on hydrophobic clusters, in addition to aggregation propensity (see also Basile et al. 2017).

Mouse-specific novel ORFs do not differ notably from ORFs created from the genic and intergenic hexamer distributions for most analysed properties (Figure 3). One exception is the length distribution, as transcriptome ORFs show much longer outliers compared to the other distributions, and hydrophobic cluster coverage. This difference is likely caused by difference in length as coverage is directly related to sequence length. More interestingly, the median aggregation propensity of unconserved transcriptome ORFs is ca. 5% lower than random ORFs generated from the intergenic as well as genic hexamer distributions. This bias in transcription likelihood could be caused by selection acting against the transcription of sequences with high aggregation propensity.

### De novo emerged ORFs resemble ORFs created by other mechanisms

We also mapped mouse ORFs onto the rat genome to determine which of the analysed ORFs represent *bona fide de novo* emerged ORFs (Table 3), i.e. evolved from intergenic regions. When analysing only the l+ORFs, 19% of the ORFs map to intergenic regions, confirming a *de novo* emergence of these ORFs. Another 15% of l+ORFs did not emerge *de novo* as they map to genic regions. The rest of the mouse-specific ORFs could not be mapped to the rat genome. It is difficult to determine the emergence mechanism of ORFs that could not be mapped to the rat genome since these ORFs either diverged drastically since the rat-mouse-split, were not present before the split, or have been lost in the rat lineage. It remains unclear if and how each of these possibilities influences the likelihood of *de novo* gene emergence. Additionally, we analysed the structural properties of the ORFs with confirmed *de novo* emergence and the ORFs with unknown or genic emergence locations (Figure S 5) to determine if different emergence mechanism lead to different structural outcomes. Most of the properties do not differ notably between the three categories, suggesting that structural and sequence properties do not depend on the emergence mechanism. Sequence length represents one exception here, as *de novo* emerged sequences are in median ~10 amino acids shorter than ORFs from either genic or unknown locations. Additionally, *de novo* ORFs have a higher fraction of sequence coverage with hydrophobic clusters. Other properties also show some deviations in medians, but these are likely caused by the multimodal distribution of the values measured for these properties.

**Table 3.**
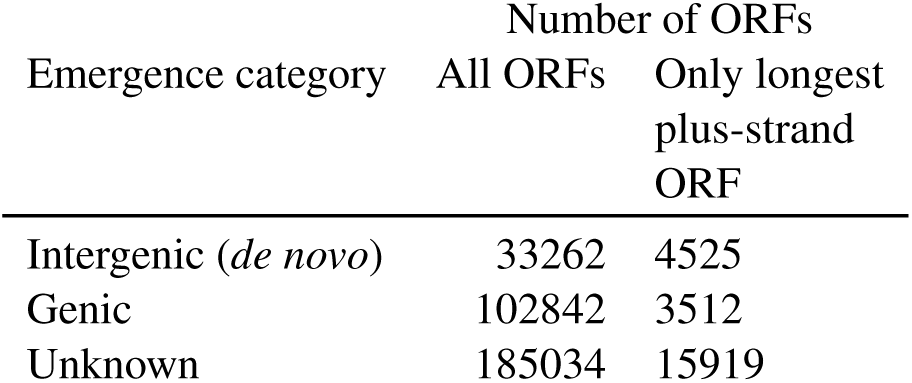
Numbers of ORFs found per location mapped to in rat. Only mouse-specific ORFs were analysed in this step. ORFs with intergenic location in rat are defined as *de novo* emerged.

| Emergence category | Number of ORFs |  |
| --- | --- | --- |
|  | All ORFs | Only longest plus-strand ORF |
| Intergenic (*de novo*) | 33262 | 4525 |
| Genic | 102842 | 3512 |
| Unknown | 185034 | 15919 |

## Conclusions

Our results suggest a new view on mechanisms and dynamics of *de novo* emergence of eukaryotic protein coding genes. A large number, certainly several thousands of newly and randomly created ORFs, are pervasively transcribed at any time and thus become exposed to selection. Most of these transcripts are rapidly lost again, probably because they do not have a beneficial function. This is in line with earlier results comparing insect genomes (Palmieri et al. 2014) and general findings on the turnover of transcription (Neme and Tautz 2016).

However, and quite unexpectedly, only hexamer score and sequence length — but not predicted disorder or aggregation propensity show — significant differences with age. The even disorder content indicates that at least those *de novo* ORFs which were not purged have similar properties to those which code for fully established proteins.

However, our findings may represent only the tip of an iceberg because ORFs coding for toxic proteins – potentially still many more than neutral ones – can not be observed, even at the very short time scale considered in our study. Still, over evolutionary very long time scales, the high loss rates diminish the number of surviving *de novo* genes to an extent that duplication or modular rearrangements (or a combination of all three mechanisms) remain the dominating process underlying the creation of new genes.

Contrary to earlier studies (Carvunis et al. 2012; Zhao et al. 2014; Wilson et al. 2017; Bornberg-Bauer, Jonathan Schmitz, et al. 2015) (see Introduction), our results indicate that, in mouse, no “structural maturation” whatsoever takes place. In that vain, the (rare) fixation of a *de novo* gene can be seen as a “frozen accident” in which the coincidentally functional state of a novel ORF becomes conserved. This is further corroborated by the finding that, contrary to an earlier study using genomic data only, different parameters for disorder prediction and random sequences (Wilson et al. 2017), randomly generated ORFs show a level of disorder that is highly similar to that of novel ORFs. Taken together, these results suggest that the nucleotide composition of the genome and the genetic code represent a predisposition that facilitates the expressibility and functional potential for spuriously occurring ORFs.

All results hold for alternative ORFs too as those can be conserved over long evolutionary time frames, albeit with a weaker correlation between age and sequence properties, suggesting that most of these ORFs do not represent emerging genes. Future studies will need to explore the function of alternative ORFs and how/if functional smORFs differ from spurious ORFs.

In an extension of earlier studies (Palmieri et al. 2014; Neme and Tautz 2013; Wilson et al. 2017) we could also locate the most likely genomic origin of many young and still mouse-specific transcribed ORFs as being intergenic, further supporting that they are bona fide *de novo* genes. Most strikingly, we find sequence properties of *de novo* ORFS being virtually indistinguishable not only from ORFs of random sequences modelled after intergenic regions, but also to those of genes which arise by divergence from existing genes. Additionally, all novel ORFs had properties different from random ORFs of random sequences modelled after the CDS nucleotide composition which shows that novel ORFs only rarely emerge from CDS, but rather from non-coding parts of genes like untranslated regions and introns. However, genes diverging from existing CDSs would not show up as young genes until they have lost all sequence similarity. Before diverging genes have lost all sequence similarity, they would still be counted as “old” genes. Therefore, no continuum of emerging diverging genes is observable. This caveat explains the high rate of duplicated and diverged genes reported in earlier studies compared to this study (Wissler et al. 2013; Donoghue et al. 2011).

Future studies will need to sample genomes and transcriptomes at even higher quality and at much shorter evolutionary gradients, e.g. at population level (Zhao et al. 2014). This will allow reconstructing the evolutionary paths of emerging ORFs in more detail and better understand the evolutionary pressures they experience.

## Materials & Methods

### Transcriptome Assembly

Mouse and other transcriptomes were assembled using the genome-guided cufflinks software (Trapnell et al. 2012), see Table 4 for data sources. For this purpose, we first aligned the trimmed reads to the genome using hisat2 with default parameters and without guide gtf (Kim et al. 2015). Trimming was performed using trimmomatic (Bolger et al. 2014), allowing for 3 seed mismatches, a palindrome threshold of 30 and a simple clip threshold of 10 (ILLUMINACLIP:3:30:10). Additionally a sliding window filtering with a window size of 4 and a required quality of 15 was employed (SLIDING-WINDOW:4:15). Afterwards, transcripts were filtered according to a length threshold that was adjusted for each library type used. After mapping, transcripts were assembled using cufflinks with default options. Here, we assembled the transcripts from each sample separately. These transcripts were then merged and filtered in the next step using cuffmerge with default options.

**Table 4.**
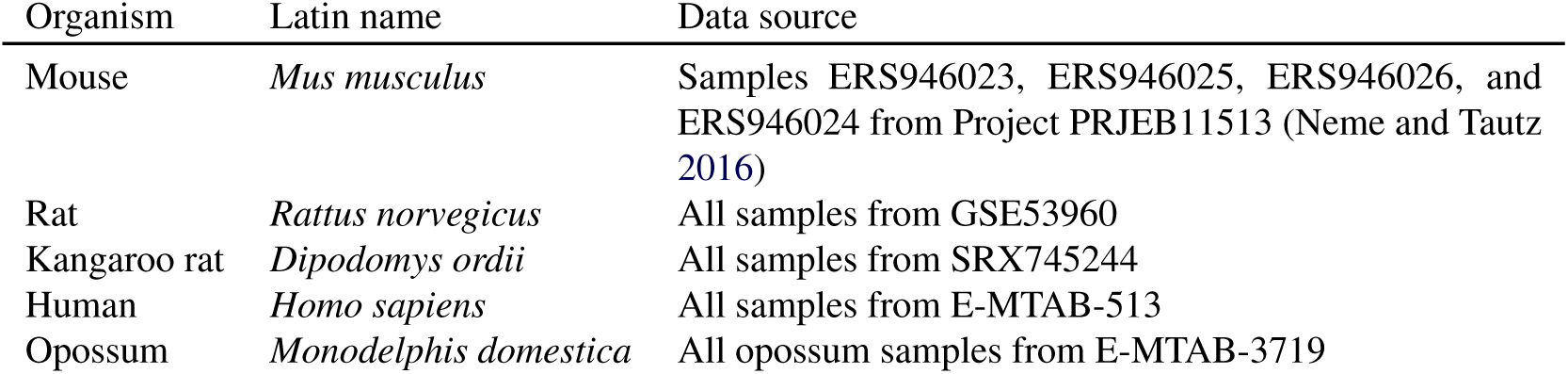
Data sources of transcriptome samples.

| Organism | Latin name | Data source |
| --- | --- | --- |
| Mouse | *Mus musculus* | Samples ERS946023, ERS946025, ERS946026, and ERS946024 from Project PRJEB11513 (Neme and Tautz 2016) |
| Rat | *Rattus norvegicus* | All samples from GSE53960 |
| Kangaroo rat | *Dipodomys ordii* | All samples from SRX745244 |
| Human | *Homo sapiens* | All samples from E-MTAB-513 |
| Opossum | *Monodelphis domestica* | All opossum samples from E-MTAB-3719 |

### ORF Prediction

ORFs were predicted using the getorf program from the EMBOSS suite (Rice et al. 2000). All sequences starting with a start codon and ending with a stop codon were considered an ORF. A minimum length of 30 amino acids was used as a threshold. ORFs were searched on the forward as well as the reverse strand. ORFs showing high similarity inside a transcriptome were filtered to only retain the longest variant. For this purpose we regarded ORFs as similar if they showed at least 95 percent identity over at least 90 percent of the shorter sequence. This step was conducted to prevent genes with copy number variation of many splice variants from skewing the distribution of sequence properties. During many analyses the data set of ORFs was restricted to only include the longest plus-strand ORF from each transcript. ORFs containing more than 25% Ns were removed from the analysis.

### ORF Age Determination

DIAMOND (Buchfink et al. 2015) and BLAST (Camacho et al. 2009) were used to search for homologous sequences in the other species’ transcriptome’s ORFs. In the analysis only including the longest plus-strand ORFs only the longest plus-strand ORFs were used, also in the four species besides mouse. Here, DIAMOND was used for a first search round. Query proteins without hits were than searched again using BLAST. An e-value cutoff of *e*^−3^ was set during these searches. The evolutionary distance to non-mouse species was set as minimum ORF age. The evolutionary distances were taken from timetree.org (“estimated” divergence times were used here).

### Sequence Property Analysis

Sequence properties were analysed using a number of programs. TANGO was used to determine aggregation propensity (Fernandez-Escamilla et al. 2004; Monsellier et al. 2007). As per the TANGO manual, stretches of at least five amino acids with an aggregation propensity higher than 5% were considered to be potentially aggregating. IUPred was used to determine intrinsic disorder in proteins (Dosztányi et al. 2005). Here, the short disorder prediction of IUPred was employed. Hydrophobic clusters of sequences were analysed using the Seq-HCA method (Faure et al. 2013). Additional basic analyses were performed using Biopython (Cock et al. 2009). No significance tests were used here, since the large number of data points in each of the categories confounds such tests and these tests do not take effect size into account.

### Random Sequence Generation

Random ORF sequences were generated by predicting ORFs on a random chromosome. This random chromosome was generated by randomly concatenating hexamers based on the hexamer frequency of different regions of the mouse genome. For this purpose we counted hexamer in intergenic, genic, and CDS regions of the genome as annotated in the GRCm38.86 version of the gtf file obtained from ensembl (Flicek et al. 2014). Here, we counted only the hexamers that were exclusive to one of the categories; no overlapping hexamers were taken into account. The mouse genome hexamer frequencies were determined using an in-house script.

### Emergence Mechanism Determination

Emergence mechanisms were determined by mapping ORFs against the rat genome using BLAT with default options. ORFs were then categorized as genic if they mapped to a gene in the rat genome or intergenic if they mapped to an intergenic region. Additionally, we classified ORFs as genic if they mapped to regions with expression evidence as per the transcriptome created earlier in this study. ORFs that did not map to the rat genome were categorized as “unknown”.

## Declarations

The datasets used and/or analysed during the current study available from the corresponding author on reasonable request. The authors declare that they have no competing interests. JS was supported by DFG grant BO2455/13-1 to EBB. All authors conceived the study. JFS performed the analysis. All authors analysed the data. JFS wrote the paper. All authors read, finalised and approved the final manuscript.

The authors would like to thank Rafik Neme for input in the initial study design. The authors would also like to thank Andreas Lange for valuable feedback on the manuscript.

**Supplementary Materials**

